# The Illumina Stranded mRNA protocol is not strongly stranded for mRNA with low U content

**DOI:** 10.64898/2026.07.09.737636

**Authors:** Olga Menshikova, Isabelle Nuez, Virginie Courtier-Orgogozo

## Abstract

The Illumina TruSeq Stranded and Illumina Stranded mRNA protocols are commonly used for strand-specific bulk RNA-seq and they typically yield >99% antisense reads. We show that these protocols can generate sense-oriented reads for transcripts with extremely low U content (<3%). Indeed, such regions can bypass the dUTP-based blockade of cDNA second strand amplification. A small number of genes are affected by this issue (three in *Drosophila melanogaster*, including the glue gene *Sgs3*, and 46 in *Mus musculus*). To prevent overestimation of expression levels, we recommend excluding sense reads for all genes.

## Introduction

RNA sequencing of populations of cells (bulk RNA-seq) has been a central method in biology and will probably remain so, given costs and flexibility in experimental design. Among the most widely used methods for bulk RNA-seq are the Illumina TruSeq™ Stranded mRNA library preparation protocol and its updated version, the Illumina Stranded protocol. Both protocols are designed to capture poly-A mRNA and preserve strand information. Because they work with a wide range of organisms and Illumina sequencing platforms, they represent a default workflow for differential gene expression experiments in many research laboratories.

When analyzing RNAseq data prepared with the Illumina TruSeq™ Stranded mRNA Illumina kit, we noticed that certain genes exhibited a significant proportion of reads opposite to the expected orientation and we investigated the origin of this puzzling observation.

## Materials and Methods

### Sample preparation

*Drosophila melanogaster* Canton S flies were raised at 25°C on a standard food medium (4 l: 83.5 g yeast, 335.0 g cornmeal, 40.0 g agar, 233.5 g saccharose, 67.0 ml Moldex, 6.0 ml propionic acid) supplemented with bromophenol blue to stage larvae (1). Dark-blue-gut, early wandering L3 female larvae were collected, washed with 1X PBS and either stored as a whole-body sample at −80°C or dissected for their pair of salivary glands. Total RNA was isolated using NucleoSpin© RNA (Macherey-Nagel). We obtained approximately 40 μL of RNA at 80-380 ng/μL for each single-larva-whole-body sample and 3-40 ng/μl for each single-pair-of-salivary-glands sample (three replicates for each condition).

### RNA sequencing

Library preparation and Illumina sequencing were performed at the Ecole normale supérieure Genomique ENS core facility (Paris, France). Messenger (polyA+) RNAs were purified from 50 ng of total RNA using oligo(dT). Libraries were prepared using the strand-specific RNA-Seq library preparation TrueSeq Stranded mRNA Prep, Ligation kit (Illumina) and were multiplexed by 30 with other samples on a P2 flow cell (Illumina). 118-bp-single-read-sequencing was performed on a NextSeq 2000 device (Illumina).

### Sequence analysis and Figures

Sequence reads were inspected before and after cleaning with FastQC version 0.12.1 and MultiQC version 1.22.2 (2). Poly N read tails and adapters were trimmed using TrimGalore! version 0.4.1 (3). Fastp version 0.23.4 (4) was used to discard reads ≤40 bases and reads with quality mean ≤30. Reads were then aligned against the *D. melanogaster* transcriptome GCF_000001215.4_Release_6_plus_ISO1_MT_rna_from_genomic_Dmel.fna or genome GCF_000001215.4_Release_6_plus_ISO1_MT_genomic.fna using HISAT2 version 2.2.1 (5) with default parameters or with kallisto version 0.50.1 (6). Samtools version 1.19.2 was used to keep only the primary mapping for each read, to extract antisense or sense reads and to extract reads mapped to specific transcripts. Read coverage was plotted with IGV-Web app version 2.4.6 (7) and Geneious Prime version 2019.2.1.

Mouse RNA-seq data was retrieved from NCBI SRA (accession SRR13657494) and processed as above, using the *M. musculus* mm10 genome. Reads mapped to specific mouse genes were retrieved based on genomic location using samtools version 1.19.2.

Sequence analyses and data visualization were performed using custom scripts written in Python version 3.12.3 and R version 4.4.2, executed in RStudio version 2024.09.1+394. Figures were prepared using LibreOffice version 24.2.5.2 (X86_64) and Inkscape version 1.4.

## Results and Discussion

### A few genes exhibit an unexpectedly high proportion of aberrantly oriented reads

In *Drosophila melanogaster* eight glue genes are specifically expressed in the salivary glands of third instar larvae and encode secreted proteins that make up an adhesive material, allowing the animal to stick to a substrate for several days (8). To identify novel glue genes, expected to be enriched in salivary glands, we prepared RNAseq from salivary glands and whole body of single *D. melanogaster* L3 larvae (3 replicates for each condition) (Fig. 1A). According to Illumina, more than 99% of the uniquely mapped reads are expected to be antisense and less than 1% sense, based on tests with human mRNA samples processed with the TruSeq Stranded mRNA (9). For our 6 samples, we were surprised to detect a higher proportion (1.9-3.1%) of sense reads than expected (Fig. 1A). Most of these aberrantly oriented reads map to the *Sgs3* glue gene transcript (Fig. 1B).

**FIGURE 1.**
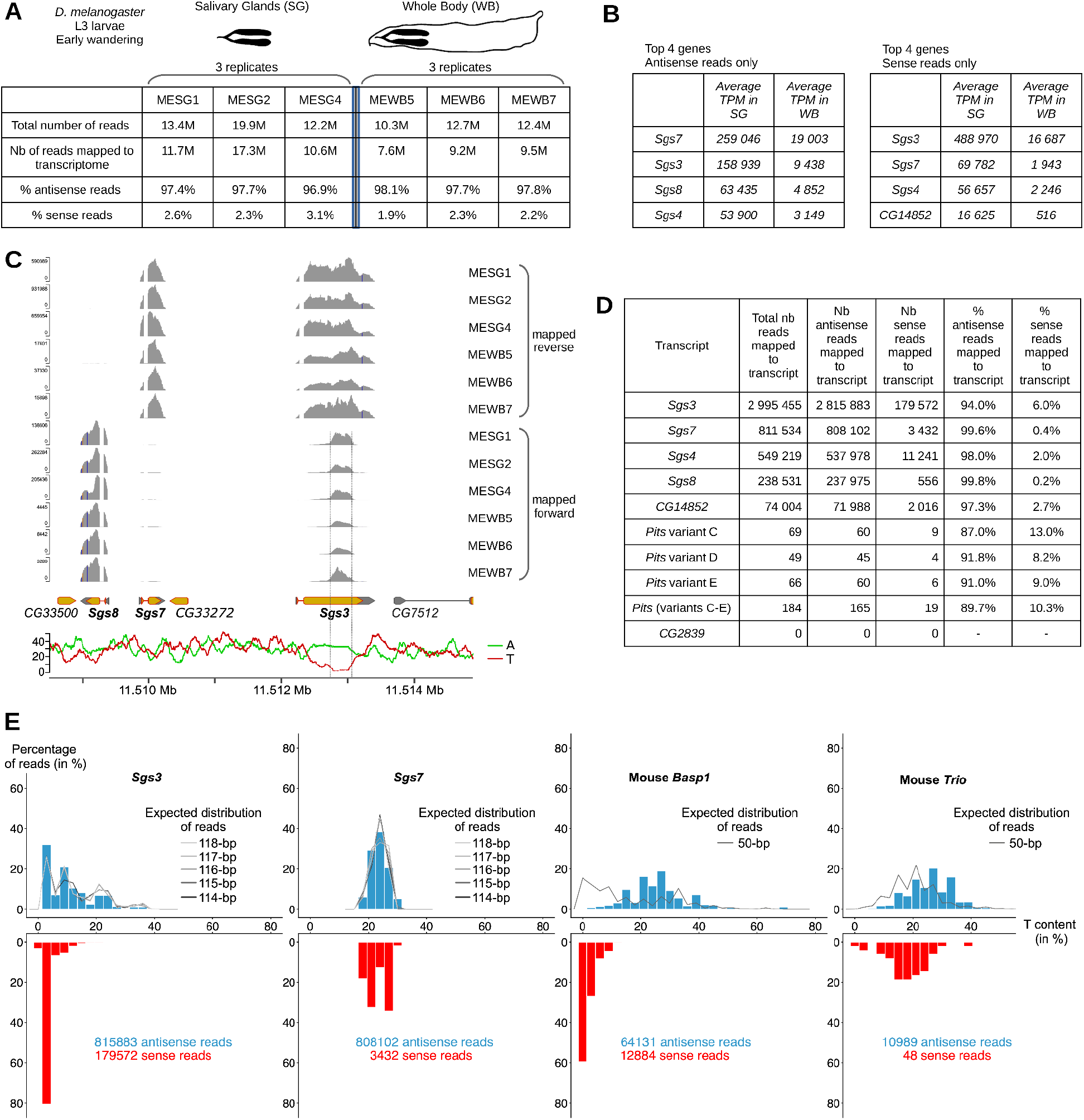
Aberrantly oriented reads correspond to low T content transcript regions. (A) Six samples (2 conditions, 3 replicates) were examined. The total number of cleaned reads and the proportion of antisense and sense reads mapped to *D. melanogaster* transcriptome were computed for each sample. (B) Top 4 genes in expression levels (TPM) in the salivary gland samples, when all reads are mapped antisense (left table) or sense (right table) relative to transcript orientation. (C) mRNA-seq read pileup tracks in the *Sgs3* genomic region for the 6 samples. Reads mapped reverse relative to genome orientation (top tracks) are distinguished from those mapped forward (bottom tracks). The %T and %A (118-bp sliding window) are indicated along the genomic region. (D) Proportion of antisense and sense reads (from the MESG1 sample) for reads mapped to specific transcripts. (E) Distribution of T content in reads from the MESG1 sample (vertical bars), when reads are antisense (top, blue) or sense (bottom, red) to transcripts *Sgs3, Sgs7, Basp1* and *Trio*. The expected distribution of T content in reads (lines) was computed by cutting in silico the transcripts into all possible fragments of a given size.

### Aberrantly oriented reads correspond to low U content transcript regions

Comparison of read coverage for antisense and sense reads mapped to the *D. melanogaster* genome in the *Sgs3* genomic region shows that the aberrantly oriented reads concentrate on a particular region of the *Sgs3* coding sequence which exhibits a low percentage of T nucleotides (Fig. 1C). The Illumina Stranded mRNA protocol relies on incorporation of dUTP instead of dTTP during second strand cDNA synthesis, and uses a DNA polymerase in a next step that does not incorporate nucleotides past dUTP. This prevents amplification of the cDNA second strand and leads ultimately to reads that are predominantly antisense (Fig. 2). We hypothesized that transcripts with low U content may bypass quenching of the cDNA second strand and result in the production of multiple sense reads for low U content regions. We compared reads mapped to *Sgs3* with those mapped to *Sgs7*, a neighboring glue gene with standard U content that is expressed in salivary glands. Sgs7 exhibits 99.6% of antisense reads and only 0.4% of sense reads. In contrast, *Sgs3* displays 94.0% of antisense reads and 6.0% of sense reads (MESG1 sample, Fig. 1D). According to the *Sgs7* transcript sequence, T content for *Sgs7* reads (when calculated along the forward gene orientation) is expected to be between 18 and 28% and this matches the observed distribution for both antisense and sense *Sgs7* reads (Fig. 1E). For *Sgs3*, the expected distribution of read T content varies from 2 to 25% and matches the observed distribution for antisense *Sgs3* reads. However antisense *Sgs3* reads are biased towards T content lower than 3% (Fig. 1E). These observations are consistent with low U content transcript regions generating reads in both orientations.

**FIGURE 2.**
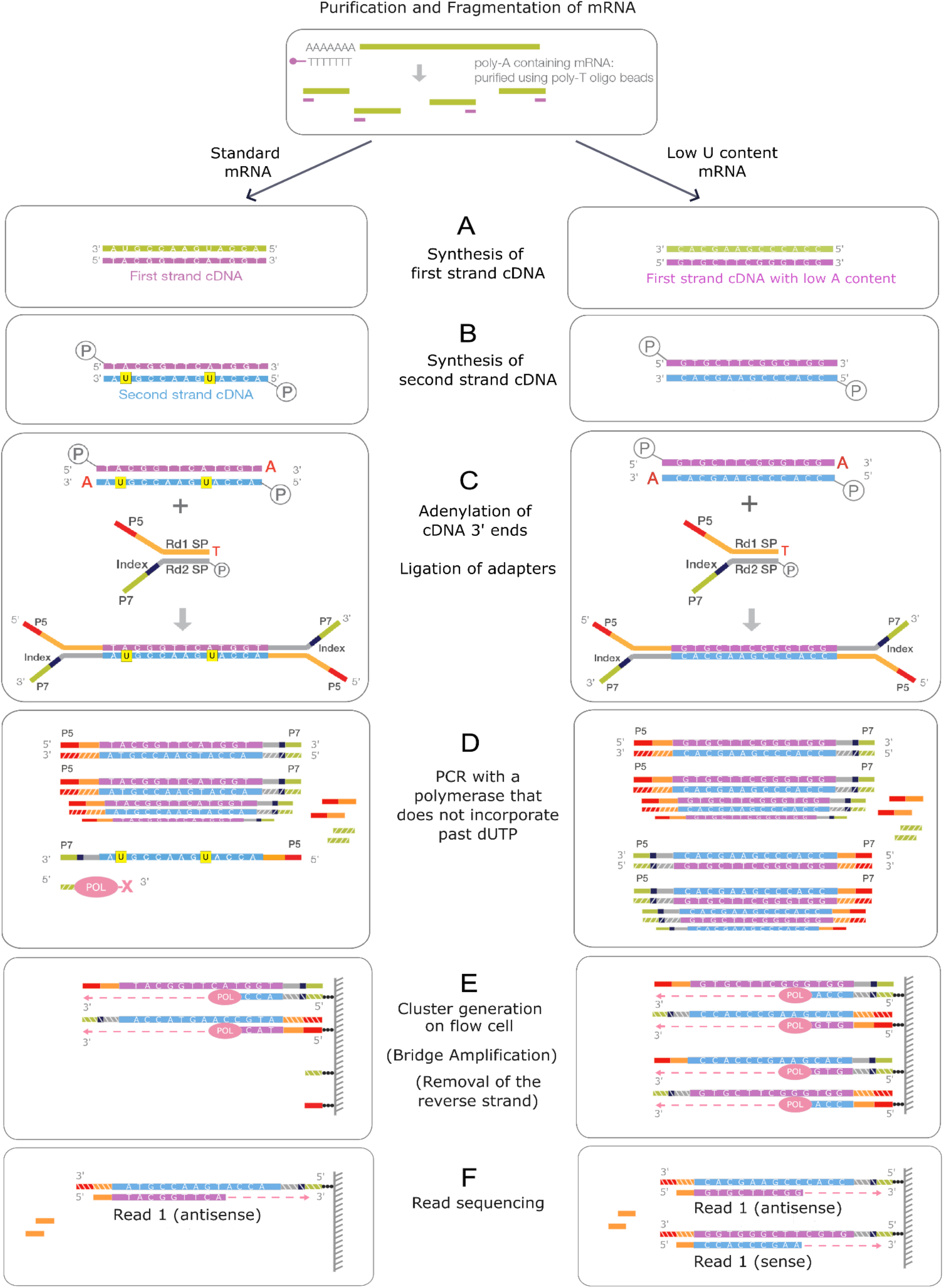
The Illumina TruSeq stranded mRNA protocol produces distinct outputs for standard mRNA molecules (left) and for low U content mRNA regions (right). Poly-A containing mRNA molecules are purified using poly-T oligos attached magnetic beads, cut into small random fragments and hybridized to a mix of random hexamers, which serve as primers for cDNA first strand synthesis (**A**). (**B**) Second strand cDNA is synthesized from a reaction mix with dUTPs instead of dTTPs. A phosphorothioate bond is added at the 5’ end. (**C**) The cDNA insert is ligated with a single A nucleotide at the 3’ end, to match the corresponding single T nucleotide at the 3’ end of the adapters. Adapters consisting of an Illumina TruSeq Universal Adapter (red and orange rectangles) and an Illumina TruSeq Index Adapter (light green and grey rectangles) are then ligated to the fragments. The Index Adapter contains a “barcode” sequence (black rectangle) for multiplex sequencing. (**D**) cDNA fragments are amplified by PCR with a DNA Polymerase that does not incorporate past dUTP, so that the second strand cDNA (light blue) is quenched during amplification (left). However, the second strand cDNA is amplified when mRNA has low U content (right). (**E**) DNA fragments are hybridized to the Illumina flow cell via the adapters. A polymerase creates a complement of the hybridized strands, after which the template is washed away. Fragments are then amplified through bridge amplification, which generates millions of clusters of fragments on the flow cell (not shown). Reverse strands are cleaved and washed off (not shown). (**F**) Forward strands, marked with specific adapters (hatched rectangles), are used for read sequencing. They generate antisense reads only (left) or both sense and antisense reads if mRNA has low U content (right). Adapted from TruSeq Stranded mRNA Reference Guide, Illumina, Document # 1000000040498 v00.

### Other low-U-content transcripts are present in Drosophila and mouse and they also lead to aberrantly oriented reads with the Illumina Stranded protocol

Examination of the *Sgs3* transcript reveals a 356-bp region containing only 10 T nucleotides (2.8% of T), to which most of the aberrantly oriented reads are mapped (Fig. 1C). We searched for regions >300bp with T content ≤2.8% in the *D. melanogaster* and *M. musculus* transcriptomes and found several transcripts: *Sgs3, Pits* and *CG2839* for *D. melanogaster*, and 46 genes (95 transcripts) for *M. musculus* (Table S1). For *Pits*, which displays low expression in our samples, we made the same observations as for *Sgs3*: we found a relatively high proportion (10.3%) of sense reads (Fig. 1D) and they were biased towards low U content.

To test whether other datasets contain aberrantly oriented reads, we analyzed a recently published mouse RNA-seq study (Illumina Stranded protocol - a more recent kit than TrueSeq, single-end 50-bp sequencing, telencephalon sample) (10). We compared *Basp1*, whose mRNA contains a 661-bp region with 2.8% T content, with the neighboring gene *Trio*, whose reads are expected to harbor between 10 and 40% of T. Results were as expected: a high proportion (16.7%) of sense reads was observed for *Basp1* but not for *Trio* (0.4%), and the *Basp1* sense reads were enriched in low T content reads (Fig. 1E).

## Discussion and Conclusion

Our analysis of Drosophila and mouse RNA-seq data shows that transcript regions with low T content (<3%) generate reads in both orientations with the TrueSeq Stranded and the Illumina Stranded mRNA kits. This affects very few protein-coding genes: only 3 (out of ~13,000) for *D. melanogaster* and 46 (out of ~21,000) for *M. musculus*. Nevertheless, this phenomenon can lead to overestimation of these genes’ expression levels. For example, the read count for *Sgs3* is increased by 20% when both sense and antisense reads are mapped (Fig. 1E). The NEBNext Ultra II Directional RNA kit also relies on U incorporation during second strand cDNA synthesis and is based on U excision (instead of the use of a DNA polymerase that does not incorporate past dUTP by Illumina kits) (11). The NEB kit should thus behave similarly. As a workaround, one possibility when preparing the libraries would be to add analogs for another nucleotide (A, C or G) alongside dUTP during the second strand cDNA synthesis step. However, the efficacy of the dUTP method is not simply due to poorer substrate tolerance of the next DNA polymerase: the stalling derives from an active uracil-recognition mechanism present in archaeal family-B polymerases (12). Other nucleotide analogs are not expected to produce such a true selective block, but rather a gradual loss of amplification. Consequently, the most effective solution is simply to exclude sense reads for all genes.

## Abbreviations

dTTP: Thymidine TriPhosphate
dUTP: DeoxyUridine TriPhosphate
Sgs: Salivary Gland Secretory
T: Thymine
TPM: Transcripts Per Million
U: Uracil

## Data Availability Statement

RNA-seq data generated in this study was deposited at NCBI Sequence Read Archive (SRA) under BioProject 1479679. Additional data that support the findings of this study are available in the Supplemental Material of this article.

## Author Contributions

V. Courtier-Orgogozo conceived and designed the research; I. Nuez prepared RNA samples, V. Courtier-Orgogozo and O. Menshikova analyzed data; V. Courtier-Orgogozo wrote the manuscript.

## Acknowledgements

We greatly thank the internship students who helped with this project: Arthur Vincens, Zoé Niemeijer and Angèle Boguais. We are grateful to Savandara Besse for help with the bioinformatics analysis and Laurent Jourdren for filtering reads. We thank Dmitri Petrov and Marie Sémon for comments on the manuscript. The GenomiqueENS core facility was supported by the France Génomique national infrastructure, funded as part of the “Investissements d’Avenir” program managed by the Agence Nationale de la Recherche (contract ANR-10-INBS-0009). This work was supported by Agence Nationale de la Recherche (ANR) under the project “ANR-24-CE13-0018-01” to V.C.-O.

## Supplemental Information

### Supplementary files

Data File S1: Fragments.csv.zip

Information about the reads mapping to the following genes of interest: *Sgs3, Sgs7, Basp1* and *Trio*. Data File used to create Fig. 1E. The csv file contains 6 columns: Gene (gene name and RNA-seq sample), Header (id number specific to each RNA-seq sample), Sequence (read sequence), T (number of T in the sequence), Sequence_Size (size of the read), Percentage_T (percentage of T in the read), Gene_Size. Each line corresponds to one read.

Data File S2: Figure1E.R

R script that uses as input Fragments.csv and creates Fig. 1E.

Data File S3: script-low-T-content-search.R

R script that uses as input a fasta file of multiple transcripts and creates as an output file a list of transcripts with regions containing few T nucleotides.

Data File S4: table_low_T_regions_Dmel_350_10.csv

Table of *D. melanogaster* transcripts that contain a region that is >350bp and whose T content is <2.86%. Output of the script script-low-T-content-search.R.

Data File S5: table_low_T_regions_Mmus_350_10.csv

Table of *M. musculus* transcripts that contain a region that is >350bp and whose T content is <2.86%. Output of the script script-low-T-content-search.R.

