## Supplementary material for "The Illumina Stranded mRNA protocol is not strongly stranded for mRNA with low U content": Data File S2: script-low-T-content-search.R

```
#Script for finding sequences with low T content
#############################################

# This script uses as input a fasta file of multiple transcripts and creates as an output file
# the list of transcripts with regions containing few T nucleotides.
# The output file is a table with 5 columns: gene name, transcript name, 
# position of beginning of the region (with a T), position of end of the region (with a T), 
# total length of the region.
#
# Two values are set in the script: 
# N_T_nucl = total number of T nucleotides in the region
# Region_length = minimal size of the region where the number of T nucleotides are counted
# Overall, the script finds regions whose size is higher than (Region_length) and whose T content is below (N_T_nucl / Region_length)

# Script written by Virginie Courtier in May 2026.


#remove.packages("BiocManager")
#install.packages("BiocManager", dependencies = TRUE)
if (!requireNamespace("BiocManager", quietly = TRUE)) install.packages("BiocManager")
BiocManager::install("Biostrings")
BiocManager::install("rtracklayer")
BiocManager::install("GenomicRanges")
library(Biostrings)
library(GenomicRanges)
library(rtracklayer)
library(dplyr)
library(stringr)

setwd("/Users/courtier/Documents/mon-labo/Droso-glue/RNAseq-STRONG-GLUE/24-11-T-content/find-transcripts-low-T")

#fasta_file <- readDNAStringSet("Sgs3-1-mel.fasta")
fasta_file <- readDNAStringSet("GCF_000001215.4_Release_6_plus_ISO1_MT_rna_Dmel.fna")
fasta_file <- readDNAStringSet("GCF_000001635.27_GRCm39_rna_Mmus.fna")


#Look for sequences with only "x" T nucleotides in a long sequence of "y" bp
#Maximum number of T nucleotides
N_T_nucl <- 10
Region_length <- 350

#Loop over the transcripts and delete intermediate data (otherwise the script creates a file that is too large) 

# Initialize an empty data frame to store the T positions
table_T_positions <- data.frame(Transcript = character(), Position = integer(), stringsAsFactors = FALSE)

# Initialize an empty data frame to store the length of regions with few T nucleotides
table_low_T_regions <- data.frame(Transcript = character(), Position_1 = integer(), Position_2 = integer(), stringsAsFactors = FALSE)

# Loop through each transcript in the FASTA file
for (i in seq_along(fasta_file)) {
  #STEP 1: Include in the first table the transcript name and position of each T within the transcript sequence
  #One line per T
  # Get the transcript name and sequence
  transcript_name <- names(fasta_file)[i]
  sequence <- fasta_file[[i]]
  #print(paste(i,": ",transcript_name))
  cat(i,": ",transcript_name," ")
  # Find positions of 'T' nucleotides
  # Splits the sequence into individual characters and then finds the indices of each 'T'
  t_positions <- which(strsplit(as.character(sequence), NULL)[[1]] == "T")
  
  # Combine transcript name with T positions and add to table_T_positions
  if (length(t_positions) > 0) {
    temp_df <- data.frame(Transcript = transcript_name, Position = t_positions, stringsAsFactors = FALSE)
    table_T_positions <- rbind(table_T_positions, temp_df)
  }

  #STEP 2: Include in the second table the transcript name, first position, last position of region with few Ts
  #One line per region
  n_final <- nrow(table_T_positions)-N_T_nucl-1
  if (n_final<1) {
    temp_df <- data.frame(Transcript = table_T_positions$Transcript[n], 
                          Position_1 = 0, 
                          Position_2 = 0,
                          stringsAsFactors = FALSE)
    table_low_T_regions <- rbind(table_low_T_regions, temp_df)}
  else
  {for (n in 1:n_final) {
      if (table_T_positions$Position[n+N_T_nucl+1]-table_T_positions$Position[n] > Region_length) {
        temp_df <- data.frame(Transcript = table_T_positions$Transcript[n], 
                              Position_1 = table_T_positions$Position[n], 
                              Position_2 = table_T_positions$Position[n+N_T_nucl+1],
                              stringsAsFactors = FALSE)
        table_low_T_regions <- rbind(table_low_T_regions, temp_df)
        cat("Y")
      }}
  }
  cat("\n")
  # Cleanup the first table
  table_T_positions <- data.frame(Transcript = character(), Position = integer(), stringsAsFactors = FALSE)
}


# View second table
View(table_low_T_regions) 
# In this table, when 0 is indicated it means that 
#the total number of Ts in the transcript is lower than (N_T_nucl + 2)

View(table(table_low_T_regions$Transcript))
#There are seven transcripts in D. mel with 10 T or less in >350bp, including the Sgs3 transcript


nrow(table_low_T_regions)

#remove lines with no transcript names and 0 values for nucleotide coordinates
table_low_T_regions <- subset(table_low_T_regions, !is.na(table_low_T_regions$Transcript))
table_low_T_regions <- subset(table_low_T_regions, table_low_T_regions$Position_1!=0)
nrow(table_low_T_regions)

table_low_T_regions$Remove <- NULL
#Keep only one line for each transcript in the table
for (i in 2:(nrow(table_low_T_regions))) {
  if (table_low_T_regions$Transcript[i]==table_low_T_regions$Transcript[i-1]) {
    table_low_T_regions$Position_1[i] <- min(table_low_T_regions$Position_1[i-1],table_low_T_regions$Position_1[i])
    table_low_T_regions$Position_2[i-1] <- max(table_low_T_regions$Position_2[i-1],table_low_T_regions$Position_2[i])
    table_low_T_regions$Remove[i-1] <- "yes"
    } else {table_low_T_regions$Remove[i-1]<- "no"}
}
#keep only lines with remove = "no"
table_low_T_regions <- subset(table_low_T_regions, table_low_T_regions$Remove=="no")
table_low_T_regions$Remove <- NULL

#compute the length (in bp) of the fragment with low U content
table_low_T_regions$Length <- table_low_T_regions$Position_2 - table_low_T_regions$Position_1 + 1

# add the name of the gene (which is the text in parentheses) as another column
table_low_T_regions$Gene <- sub(".*\\(([^()]*)\\).*", "\\1", table_low_T_regions$Transcript)

#place the column "Gene" as the first column in the table
#sort table according to gene names
table_low_T_regions <- table_low_T_regions %>% relocate(Gene)
table_low_T_regions <- table_low_T_regions %>% arrange(Gene)

write.csv(table_low_T_regions, "table_low_T_regions_Dmel_350_10.csv", row.names = FALSE)
write.csv(table_low_T_regions, "table_low_T_regions_Mmus_350_10.csv", row.names = FALSE)
View(table(table_low_T_regions$Gene))
#There are 46 genes with low T content regions in the Mus musculus transcriptome.


table_low_T_regions <- read.csv("table_low_T_regions_Dmel_350_10.csv", sep=",", fileEncoding = "UTF-8")
table_low_T_regions <- read.csv("table_low_T_regions_Mmus_350_10.csv", sep=",", fileEncoding = "UTF-8")

#########################
#Find non coding transcripts that might be been misidentified due to sense reads aberrantly mapped to the
# transcripts with low T content regions


#First look for transcripts that overlap with the transcripts identified in table_low_T_regions.

#DID NOT FIND ANYTHING

#extract all the characters up to the first space = extract the NM_ accession number
table_low_T_regions$Transcript_ID <- str_extract(table_low_T_regions$Transcript, "^[^ ]+")
#prepare the list of genes for which we want to search for an overlapping transcript in the opposite strand
transcripts_of_interest <- names(table(table_low_T_regions$Transcript_ID))
genes_of_interest  <- names(table(table_low_T_regions$Gene))

#include Pde4/dnc gene as a control because it contains several genes nested within one of its intron
genes_of_interest <- c(genes_of_interest,"dnc")

annotations <- import("GCF_000001215.4_Release_6_plus_ISO1_MT_genomic.gff")
annotations <- import("GCF_000001635.27_GRCm39_genomic.gff")
transcripts <- annotations[annotations$type %in% c("transcript", "mRNA")]

#identify the gene name field
possible_cols <- c("gene_name", "gene", "Name", "gene_id")
gene_col <- possible_cols[possible_cols %in% names(mcols(transcripts))][1]
gene_names <- mcols(transcripts)[[gene_col]]
#the field is "gene"

#transcripts$gene_names

head(gene_names)
head(genes_of_interest)
head(transcripts_of_interest)
View(intersect(gene_names, genes_of_interest))
#there are 33 intersected names
View(table(genes_of_interest))
nrow(table(genes_of_interest))
#there are 46 genes of interest, in which to search for opposite strand transcripts
#so 13 genes are MISSING IN THE SEARCH _ FIND PROBLEM

#the three genes of D.mel are present in the gff annotation file

query_tx <- transcripts[gene_names %in% genes_of_interest]

#find overlaping genes
#hits <- findOverlaps(query_tx, transcripts, ignore.strand = FALSE)
#not good because the transcrpts should overlap
#I prefer to search for transcripts that could be nested within introns as well
gene_regions <- reduce(query_tx, ignore.strand = TRUE)

hits <- findOverlaps(transcripts, gene_regions, ignore.strand = TRUE)

tx_hits <- transcripts[queryHits(hits)]
region_hits <- gene_regions[subjectHits(hits)]

# keep only different genes
tx_gene <- mcols(tx_hits)[[gene_col]]
query_gene_set <- unique(genes_of_interest)
query_gene_set

result <- data.frame(
  chr = as.character(seqnames(tx_hits)),
  start = start(tx_hits),
  end = end(tx_hits),
  strand = as.character(strand(tx_hits)),
  gene = tx_gene,
  stringsAsFactors = FALSE
)

View(result)
write.csv(result, "table_rev-and-for-genes-low_T_regions_Mmus_350_10.csv", row.names = FALSE)
#I inspected all of these transcripts manually and most are nested within introns.
#I did not find any transcripts that would be abberantly annotated due to sense reads via Illumina Stranded protocol.
```
