## Supplementary material for "The Illumina Stranded mRNA protocol is not strongly stranded for mRNA with low U content": Data File S3: Figure1E.R

```
#Script for creating Fig. 1E of paper "The Illumina Stranded mRNA protocol is not strongly stranded for mRNA with low U content"
#############################################


#This scripts builds a ggplot of the distribution of T content for sense and antisense reads 
# for the following genes: D. melanogaster Sgs3, D. melanogaster Sgs7, M. musculus Basp1 and M. musculus Pits.

#This script was written by Angèle Boguais in May-June 2026.

#The input file for this script is Fragments.csv (the input file should be in the working directory).

#The plot are mirror graphs that shows the number in log scale of reads per percentage of T.
#The abscissa indicates the percentage of T.
#The ordinate indicates the number of reads. On top of the abscissa are the antisense reads and at the bottoms are the sense reads
# The plot contains both a Histograms and a frequency polygon.
# The frequency polygon represents the distribution of theoretical reads.
# For the theoretical reads, the number of sense reads is always 0, because by the Illumina protocol, there should be near 0 reads sense. 
# The histogram represents the experimental data
# The plots are going to be build depending on the contents of the Fragments.csv file
# There should be  at least 3 columns inside the Fragments.csv file (the Fragments.csv file actually contains 7 columns) :
# Gene (gene name and RNA-seq sample) 
# Sequence_Size (size of the read)
# Percentage_T (percentage of T in the read)
# The genes should be named appropriately depending of the genes :
# Sgs3 :  
# Sgs3-RA-mel : for the theoretical reads
# MESG1_Sgs3_sense or MEWB5_Sgs3_sense : for the experimental reads of the sample MESG1 or MEWB5
# MESG1_Sgs3_antisense or MEWB5_Sgs3_antisense : for the experimental reads of the sample MESG1 or MEWB5
# Sgs7 :  
# Sgs7-RA-mel : for the theoretical reads
# MESG1_Sgs7_sense or MEWB5_Sgs7_sense : for the experimental reads of the sample MESG1 or MEWB5
# MESG1_Sgs7_antisense or MEWB5_Sgs7_antisense : for the experimental reads of the sample MESG1 or MEWB5
#  Basp1 :
#  Basp1-RA-mus : for the theoretical reads
#  Basp1_reverse : sens reads
#  Basp1_forward : antisens reads
# Pits : 
# PitsC-RA-mel
# PitsE-RA-mel
# PitsD-RA-mel
# MESG1_Pits_sense
# MESG1_Pits_antisense

setwd("/Users/courtier/Documents/mon-labo/membres/stagiaires/26-05-Angele-BOGUAIS/scripts-Angele")

data <- read.csv("Fragments.csv", header = TRUE, sep = ",")

data$Sequence_Size<- factor(data$Sequence_Size, ordered= TRUE)

data<-data.frame(data)
# install.packages("ggplot2")
# install.packages("patchwork")

library(ggplot2)
library(patchwork)


#reads theoretical 
sgs3<- data[data$Gene == 'Sgs3-RA-mel'& (data$Sequence_Size == 114 |data$Sequence_Size == 115|data$Sequence_Size == 116 |data$Sequence_Size == 117 |data$Sequence_Size == 118),]
# experimental sens of MESG1
MESG1sgs3sens <- data[data$Gene == 'MESG1_Sgs3_sense',]
# experimental antisense readsof MESG1
MESG1sgs3antisens <- data[data$Gene == 'MESG1_Sgs3_antisense',]
# experimental antisense reads of MEWB5
MEWB5sgs3antisens <- data[data$Gene == 'MEWB5_Sgs3_antisense',]
# experimental sens reads of MEWB5
MEWB5sgs3sens <- data[data$Gene == 'MEWB5_Sgs3_sense',]


#reads theoretical 
sgs7<- data[data$Gene == 'Sgs7-RA-mel'& (data$Sequence_Size == 114 |data$Sequence_Size == 115|data$Sequence_Size == 116 |data$Sequence_Size == 117 |data$Sequence_Size == 118) ,]
# experimental sens reads of MESG1
MESG1sgs7sens <- data[data$Gene == 'MESG1_Sgs7_sense',]
# experimental antisense reads of MESG1
MESG1sgs7antisens <- data[data$Gene == 'MESG1_Sgs7_antisense',]
# experimental antisense reads of MEWB5
MEWB5sgs7antisens <- data[data$Gene == 'MEWB5_Sgs7_antisense',]
# experimental sens reads of MEWB5
MEWB5sgs7sens <- data[data$Gene == 'MEWB5_Sgs7_sense',]

#reads theoretical 
Basp1<- data[data$Gene == 'Basp1-RA-mus'& data$Sequence_Size == 50,]
# experimental sens reads of 50 pb
Basp1sens <- data[data$Gene == 'Basp1_reverse',]
# experimental antisense reads of 50 pb
Basp1antisens <- data[data$Gene == 'Basp1_forward',]

#reads theoretical 
Trio<- data[data$Gene == 'Trio-RA-mus' & data$Sequence_Size == 50, ]
# experimental sens reads of 50 pb
Triosens <- data[data$Gene == 'Trio_reverse',]
# experimental antisense reads of 50 pb
Trioantisens <- data[data$Gene == 'Trio_forward',]

#reads theoretical 
PitsC<- data[data$Gene == 'PitsC-RA-mel' & (data$Sequence_Size == 118) ,]
PitsE<- data[data$Gene == 'PitsE-RA-mel' & (data$Sequence_Size == 118) ,]
PitsD<- data[data$Gene == 'PitsD-RA-mel' & (data$Sequence_Size == 118) ,]
# experimental sens reads
Pitssens <- data[data$Gene == 'MESG1_Pits_sense',]
# experimental antisense reads 
Pitsantisens <- data[data$Gene == 'MESG1_Pits_antisense',]

oranges   <- colorRampPalette(c("#ffd6ad", "#e6550d"))(length(sort(unique(sgs3$Sequence_Size))))

MESG1Sgs3Top <- ggplot() +   
  geom_histogram(data = MESG1sgs3antisens, binwidth = 3, aes(x = Percentage_T, y=after_stat(count / sum(count) * 100), fill = "Experimental reads antisens"),color="white") +
  geom_freqpoly(data = sgs3, binwidth = 3, aes(x = Percentage_T, y=after_stat(count / tapply(count, group, sum)[group] * 100), group = Sequence_Size, colour = factor(Sequence_Size))) +
  coord_cartesian(xlim = c(0, 80), ylim = c(0, 83)) + 
  scale_y_continuous(name="Percentage of reads", labels =abs )+
  scale_fill_manual(values = c("Experimental reads antisens" = "#3399cc")) +
 scale_color_manual(values = gray.colors(length(unique(sgs3$Sequence_Size)), start = 0.2, end = 0.8),name = "Reads size")+
  theme_classic() +
  theme(axis.title.x = element_blank(),axis.text.x = element_blank(),axis.title.y = element_blank(),legend.position = "none")+ 
  ggtitle("Sgs3 MESG1") +
  labs(color='Reads size',fill ='') +
  #Descriptions on the figure
  annotate("text", x = 25, y = 50, label = paste("Antisens :", nrow(MESG1sgs3antisens)), color = "#3399cc",size = 7) +
  theme( legend.title = element_text(size = 10),
         legend.text = element_text( size = 10),
         axis.text.y = element_text(size = 18))

MESG1Sgs3Bottom <- ggplot() +   
  geom_histogram(data = MESG1sgs3sens, binwidth = 3,
                 aes(x = Percentage_T,  y=after_stat(count / sum(count) * 100), fill = "Experimental reads sens"),color="white") +
  scale_y_reverse(name="Percentage of reads") + 
  coord_cartesian(xlim = c(0, 80), ylim = c(83, 0)) + 
  scale_fill_manual(values = c("Experimental reads sens" = "#ff0000")) +
  #geom_freqpoly(data = sgs3, binwidth = 3,aes(x=Percentage_T,y=after_stat(count / tapply(count, group, sum)[group] * -100)*0, group=Sequence_Size,colour = factor(Sequence_Size)))+
  theme_classic() +
 #scale_color_manual(values = gray.colors(length(unique(sgs3$Sequence_Size)), start = 0.2, end = 0.8),name = "Reads size")+

  scale_x_continuous(name = "Percentage of T",position = "top") +
  theme(axis.text = element_text(size = 18),axis.title.y = element_blank(),axis.title.x = element_blank() )+
  labs(color='Reads size',fill ='') +
  #Descriptions on the figure
  annotate("text", x = 25, y = 50, label = paste("Sens :", nrow(MESG1sgs3sens)),color = "#ff0000", size = 7) +
  theme( legend.title = element_text(size = 10),
         legend.text = element_text( size = 10),legend.position = "none",
         axis.text.y = element_text(size = 18))

MESG1Sgs3Plot <- (MESG1Sgs3Top / MESG1Sgs3Bottom)  +
  plot_layout(heights = c(1, 1)) & 
  theme(plot.margin = margin(t = 0, b = 0))

MESG1Sgs7Top <- ggplot() +   
  geom_histogram(data = MESG1sgs7antisens, binwidth = 3, aes(x = Percentage_T, y=after_stat(count / sum(count) * 100), fill = "Experimental reads antisens"),color="white") +
  geom_freqpoly(data = sgs7, binwidth = 3, aes(x = Percentage_T, y=after_stat(count / tapply(count, group, sum)[group] * 100), group = Sequence_Size, colour = factor(Sequence_Size))) +
  coord_cartesian(xlim = c(0, 80), ylim = c(0, 83)) + 
  scale_y_continuous( labels =abs )+
  scale_fill_manual(values = c("Experimental reads antisens" = "#3399cc")) +
scale_color_manual(values = gray.colors(length(unique(sgs7$Sequence_Size)), start = 0.2, end = 0.8),name = "Reads size")+
  theme_classic() +
  theme(axis.title.y = element_blank(),axis.title.x = element_blank(),axis.text.x = element_blank(),legend.position = "none")+ 
  ggtitle("Sgs7 MESG1") +
  labs(color='Reads size',fill ='') +
  #Descriptions on the figure
  annotate("text", x = 25, y = 50, label = paste("Antisens :", nrow(MESG1sgs7antisens)),color = "#3399cc", size = 7) +
  theme( legend.title = element_text(size = 10),
         legend.text = element_text( size = 10),
         axis.text.y = element_text(size = 18))

MESG1Sgs7Bottom <- ggplot() +   
  geom_histogram(data = MESG1sgs7sens, binwidth = 3, aes(x = Percentage_T,  y=after_stat(count / sum(count) * 100), fill = "Experimental reads sens"),color="white") +
  #geom_freqpoly(data = sgs7, binwidth = 3,aes(x=Percentage_T,y=after_stat(count / tapply(count, group, sum)[group] * -100)*0, group=Sequence_Size,colour = factor(Sequence_Size)))+
  scale_y_reverse() + 
  coord_cartesian(xlim = c(0, 80), ylim = c(83, 0)) + 
  scale_fill_manual(values = c("Experimental reads sens" = "#ff0000")) +
   theme_classic() +
 #scale_color_manual(values = gray.colors(length(unique(sgs7$Sequence_Size)), start = 0.2, end = 0.8),name = "Reads size")+

  scale_x_continuous(name = "Percentage of T",position = "top") +
  theme(axis.title.y = element_blank(),axis.title.x = element_blank(),axis.text = element_text(size = 18),legend.position = "none")+
  labs(color='Reads size',fill ='') +
  #Descriptions on the figure
  annotate("text", x = 25, y = 50, label = paste("Sens :", nrow(MESG1sgs7sens)), color = "#ff0000",size = 7) +
  theme( legend.title = element_text(size = 10),
         legend.text = element_text( size = 10),
         axis.text.y = element_text(size = 18))

MESG1Sgs7Plot <- (MESG1Sgs7Top / MESG1Sgs7Bottom) +
  plot_layout(heights = c(1, 1)) & 
  theme(plot.margin = margin(t = 0, b = 0))

SgsPlot <- (MESG1Sgs3Plot | MESG1Sgs7Plot)   
  #+plot_layout(guides = 'collect') 

MEWB5Sgs3Top <- ggplot() +   
  geom_histogram(data = MEWB5sgs3antisens, binwidth = 3, aes(x = Percentage_T, y=after_stat(count / sum(count) * 100), fill = "Experimental reads antisens"),color="white") +
  geom_freqpoly(data = sgs3, binwidth = 3, aes(x = Percentage_T, y=after_stat(count / tapply(count, group, sum)[group] * 100), group = Sequence_Size, colour = factor(Sequence_Size))) +
  coord_cartesian(xlim = c(0, 80), ylim = c(0, 83)) + 
  scale_y_continuous(name="Percentage of reads", labels =abs )+
  scale_fill_manual(values = c("Experimental reads antisens" = "#3399cc")) +
 scale_color_manual(values = gray.colors(length(unique(sgs3$Sequence_Size)), start = 0.2, end = 0.8),name = "Reads size")+

  theme_classic() +
  theme(axis.title.y = element_blank(),axis.title.x = element_blank(),axis.text.x = element_blank()) + 
  ggtitle("Sgs3 MEWB5") +
  labs(color='Reads size',fill ='') +
  #Descriptions on the figure
  annotate("text", x = 25, y = 50, label = paste("Antisens :", nrow(MEWB5sgs3antisens)), color ="#3399cc", size = 7) +
  theme( legend.title = element_text(size = 10),
         legend.text = element_text( size = 10),
         axis.text.y = element_text(size = 18))

MEWB5Sgs3Bottom <- ggplot() +   
  geom_histogram(data = MEWB5sgs3sens, binwidth = 3,
                 aes(x = Percentage_T,  y=after_stat(count / sum(count) * 100), fill = "Experimental reads sens"),color="white") +
  scale_y_reverse(name="Percentage of reads") + 
  coord_cartesian(xlim = c(0, 80), ylim = c(83, 0)) + 
  scale_fill_manual(values = c("Experimental reads sens" = "#ff0000")) +
  #geom_freqpoly(data = sgs3, binwidth = 3,aes(x=Percentage_T,y=after_stat(count / tapply(count, group, sum)[group] * -100)*0, group=Sequence_Size,colour = factor(Sequence_Size)))+
  theme_classic() +
 #scale_color_manual(values = gray.colors(length(unique(sgs3$Sequence_Size)), start = 0.2, end = 0.8),name = "Reads size")+

  scale_x_continuous(name = "Percentage of T",position = "top") +
  labs(color='Reads size',fill ='') +
  #Descriptions on the figure  
  annotate("text", x = 25, y = 50, color = "#ff0000",label = paste("Sens :", nrow(MEWB5sgs3sens)), size = 7) +
  theme( legend.title = element_text(size = 10),
         legend.text = element_text( size = 10),axis.title.y = element_blank(),axis.title.x = element_blank(),legend.position = "none",
         axis.text.y = element_text(size = 18))

MEWB5Sgs3Plot <- (MEWB5Sgs3Top / MEWB5Sgs3Bottom)+
  plot_layout(heights = c(1, 1)) & 
  theme(plot.margin = margin(t = 0, b = 0)) 

MEWB5Sgs7Top <- ggplot() +   
  geom_histogram(data = MEWB5sgs7antisens, binwidth = 3, aes(x = Percentage_T, y=after_stat(count / sum(count) * 100), fill = "Experimental reads antisens"),color="white") +
  geom_freqpoly(data = sgs7, binwidth = 3, aes(x = Percentage_T, y=after_stat(count / tapply(count, group, sum)[group] * 100), group = Sequence_Size, colour = factor(Sequence_Size))) +
  coord_cartesian(xlim = c(0, 80), ylim = c(0, 83)) + 
  scale_y_continuous( labels =abs )+
  scale_fill_manual(values = c("Experimental reads antisens" = "#3399cc")) +
 scale_color_manual(values = gray.colors(length(unique(sgs7$Sequence_Size)), start = 0.2, end = 0.8),name = "Reads size")+

  theme_classic() +
  theme(axis.title.x = element_blank(),axis.title.y = element_blank(),axis.text.x = element_blank()) + 
  ggtitle("Sgs7 MEWB5") +
  labs(color='Reads size',fill ='') +
  #Descriptions on the figure
  annotate("text", x = 25, y = 50, label = paste("Antisens :", nrow(MEWB5sgs7antisens)), color = "#3399cc",size = 7) +
  theme( legend.title = element_text(size = 10),
         legend.text = element_text( size = 10),axis.title.y = element_blank(),axis.title.x = element_blank(),legend.position = "none",
         axis.text.y = element_text(size = 18))

MEWB5Sgs7Bottom <- ggplot() +   
  geom_histogram(data = MEWB5sgs7sens, binwidth = 3, aes(x = Percentage_T,  y=after_stat(count / sum(count) * 100), fill = "Experimental reads sens"),color="white") +
  #geom_freqpoly(data = sgs7, binwidth = 3,aes(x=Percentage_T,y=after_stat(count / tapply(count, group, sum)[group] * -100)*0, group=Sequence_Size,colour = factor(Sequence_Size)))+
  scale_y_reverse() + 
  coord_cartesian(xlim = c(0, 80), ylim = c(83, 0)) + 
  scale_fill_manual(values = c("Experimental reads sens" = "#ff0000")) +
  theme_classic() +
 #scale_color_manual(values = gray.colors(length(unique(sgs7$Sequence_Size)), start = 0.2, end = 0.8),name = "Reads size")+

  scale_x_continuous(name = "Percentage of T",position = "top") +
  labs(color='Reads size',fill ='') +
  #Descriptions on the figure
  annotate("text", x = 25, y = 50, label = paste("Sens :", nrow(MEWB5sgs7sens)), color = "#ff0000",size = 7) +
  theme( legend.title = element_text(size = 10),legend.position = "none",
         legend.text = element_text( size = 10), axis.title.y = element_blank(),axis.title.x = element_blank(),
         axis.text.y = element_text(size = 18))

MEWB5Sgs7Plot <- (MEWB5Sgs7Top / MEWB5Sgs7Bottom)+
  plot_layout(heights = c(1, 1)) & 
  theme(plot.margin = margin(t = 0, b = 0)) 

MEWB5SgsPlot <- (MEWB5Sgs3Plot | MEWB5Sgs7Plot)
# +  plot_layout(guides = 'collect')

Basp1Top <- ggplot() +   
  geom_histogram(data = Basp1antisens, binwidth = 3, aes(x = Percentage_T, y=after_stat(count / sum(count) * 100), fill = "Experimental reads antisens"),color="white") +
  geom_freqpoly(data = Basp1, binwidth = 3, aes(x = Percentage_T, y=after_stat(count / tapply(count, group, sum)[group] * 100), group = Sequence_Size, colour = factor(Sequence_Size))) +
  coord_cartesian(xlim = c(0, 80), ylim = c(0, 83)) + 
  scale_y_continuous(name="Percentage of reads", labels =abs )+
  scale_fill_manual(values = c("Experimental reads antisens" = "#3399cc")) +
  scale_color_manual(values = c( "#636362")) +
  theme_classic() +
  theme(axis.title.x = element_blank(),axis.text.x = element_blank()) + 
  ggtitle("Basp1") +
  labs(color='Reads size',fill ='') +
  #Descriptions on the figure
  annotate("text", x = 25, y = 50, label = paste("Antisens :", nrow(Basp1antisens)),color = "#3399cc", size = 7) +
  theme( legend.title = element_text(size = 10),  legend.position = "none", 
         legend.text = element_text( size = 10), axis.title.y = element_blank(),axis.title.x = element_blank(),
         axis.text.y = element_text(size = 18))

Basp1Bottom <- ggplot() +   
  geom_histogram(data = Basp1sens, binwidth = 3,
                 aes(x = Percentage_T,  y=after_stat(count / sum(count) * 100), fill = "Experimental reads sens"),color="white") +
  scale_y_reverse(name="Percentage of reads") + 
  coord_cartesian(xlim = c(0, 80), ylim = c(83, 0)) + 
  scale_fill_manual(values = c("Experimental reads sens" = "#ff0000")) +
  #geom_freqpoly(data =Basp1, binwidth = 3,aes(x=Percentage_T,y=after_stat(count / tapply(count, group, sum)[group] * -100)*0, group=Sequence_Size,colour = factor(Sequence_Size)))+
  theme_classic() + 
  #scale_color_manual(values = c( "#636362")) +
  scale_x_continuous(name = "Percentage of T",position = "top") +
  labs(color='Reads size',fill ='') +
  #Descriptions on the figure
  annotate("text", x = 20, y = 50, label = paste("Sens :", nrow(Basp1sens)),color = "#ff0000", size = 7) +
  theme( legend.title = element_text(size = 10), axis.title.y = element_blank(),axis.title.x = element_blank(),
         legend.text = element_text( size = 10),legend.position = "none", axis.text = element_text(size = 18),
         axis.text.y = element_text(size = 18))

Basp1Plot <- (Basp1Top / Basp1Bottom)+
  plot_layout(heights = c(1, 1))& theme(plot.margin = margin(t = 0, b = 0)) 

TrioTop <- ggplot() +   
  geom_histogram(data = Trioantisens, binwidth = 3, aes(x = Percentage_T, y=after_stat(count / sum(count) * 100), fill = "Experimental reads antisens"),color="white") +
  geom_freqpoly(data = Trio, binwidth = 3, aes(x = Percentage_T, y=after_stat(count / tapply(count, group, sum)[group] * 100), group = Sequence_Size, colour = factor(Sequence_Size))) +
  coord_cartesian(xlim = c(0, 80), ylim = c(0, 83)) + 
  scale_y_continuous(name="Percentage of reads", labels =abs )+
  scale_fill_manual(values = c("Experimental reads antisens" = "#3399cc")) +
  
  scale_color_manual(values = c( "#636362")) +
  theme_classic() +
  theme(axis.title.x = element_blank(),axis.text.x = element_blank()) + 
  ggtitle("Trio") +
  labs(color='Reads size',fill ='') +
  #Descriptions on the figure
  annotate("text", x = 25, y = 50, label = paste("Antisens :", nrow(Trioantisens)), color = "#3399cc",size = 7) +
  theme( legend.title = element_text(size = 10),legend.position = "none",
         legend.text = element_text( size = 10),axis.title.y = element_blank(),axis.title.x = element_blank(),
         axis.text.y = element_text(size = 18))

TrioBottom <- ggplot() +   
  geom_histogram(data = Triosens, binwidth = 3, aes(x = Percentage_T,  y=after_stat(count / sum(count) * 100), fill = "Experimental reads sens"),color="white") +
  #geom_freqpoly(data = Trio, binwidth = 3,aes(x=Percentage_T,y=after_stat(count / tapply(count, group, sum)[group] * -100)*0, group=Sequence_Size,colour = factor(Sequence_Size)))+
  scale_y_reverse(name="Percentage of reads") + 
  coord_cartesian(xlim = c(0, 80), ylim = c(83, 0)) + 
  scale_fill_manual(values = c("Experimental reads sens" = "#ff0000")) +
  theme_classic() +
  #scale_color_manual(values = c( "#636362")) +
  scale_x_continuous(name = "Percentage of T",position = "top") +
  labs(color='Reads size',fill ='') +
  #Descriptions on the figure
  annotate("text", x = 25, y = 50, label = paste("Sens :", nrow(Triosens)), color = "#ff0000",size = 7) +
  theme( legend.title = element_text(size = 10),axis.title.y = element_blank(),axis.title.x = element_blank(),
         legend.text = element_text( size = 10), legend.position = "none", axis.text = element_text(size = 18),
         axis.text.y = element_text(size = 18))

TrioPlot <- (TrioTop / TrioBottom)+
  plot_layout(heights = c(1, 1)) & 
  theme(plot.margin = margin(t = 0, b = 0)) 

Basp1TrioPlot <- (Basp1Plot | TrioPlot) +  
  plot_layout(guides = 'collect') 


IlluminaPlot <-
  MESG1Sgs3Plot |
  MESG1Sgs7Plot |
  Basp1Plot |
  TrioPlot 

IlluminaPlot

ggsave(
  "IlluminaPlot.pdf",
  IlluminaPlot,
  width = 60,
  height = 22,
  units = "cm",
)

sizeC <- sort(unique(PitsC$Sequence_Size))
purples <- colorRampPalette(c("#f39cff", "#931ca3"))(length(sizeC))
names(purples) <- paste("C", sizeC)
sizeD<- sort(unique(PitsD$Sequence_Size))
greens <- colorRampPalette(c("#a1d99b", "#006d2c"))(length(sizeD))
names(greens) <- paste("D", sizeD)
sizeE<- sort(unique(PitsE$Sequence_Size))
oranges2   <- colorRampPalette(c("#ffd6ad", "#e6550d"))(length(sizeE))
names(oranges2) <- paste("E", sizeE)
shadings <- c(purples, greens, oranges2)


PitsTop <- ggplot() +   
  geom_histogram(data = Pitsantisens, binwidth = 3, aes(x = Percentage_T, y=after_stat(count / sum(count) * 100), fill = "Experimental reads antisens"),color="white") +
  geom_freqpoly(data = PitsE, binwidth = 3,aes(x=Percentage_T,y=after_stat(count / tapply(count, group, sum)[group]) * 100,group=Sequence_Size, color = paste("E", Sequence_Size)))+ 
  geom_freqpoly(data = PitsD, binwidth = 3,aes(x=Percentage_T,y=after_stat(count / tapply(count, group, sum)[group]) * 100,group=Sequence_Size, color = paste("D", Sequence_Size)))+  
  geom_freqpoly(data = PitsC, binwidth = 3,aes(x=Percentage_T,y=after_stat(count / tapply(count, group, sum)[group]) * 100, group = Sequence_Size, color = paste("C", Sequence_Size)))+

  coord_cartesian(xlim = c(0, 80), ylim = c(0, 83)) + 
  scale_y_continuous(name="Percentage of reads", labels =abs )+
  scale_fill_manual(values = c("Experimental reads antisens" = "#3399cc")) +
  #scale_color_grey()+
  theme_classic() +
  theme(axis.title.x = element_blank(),axis.text.x = element_blank(),legend.position = "none") + 
  ggtitle("Pits") +
  labs(color='Reads size',fill ='') +
  #Descriptions on the figure  
  annotate("text", x = 25, y = 50, color = "#3399cc",label = paste("Antisens :", nrow(Pitsantisens)), size = 7)+
  scale_color_manual(values = shadings) +
  theme( legend.title = element_text(size = 10),legend.position = "none",
         legend.text = element_text( size = 10), axis.title.y = element_blank(),axis.title.x = element_blank(),
         axis.text.y = element_text(size = 18))

PitsBottom <- ggplot() +   
  geom_histogram(data = Pitssens, binwidth = 3,
                 aes(x = Percentage_T,  y=after_stat(count / sum(count) * 100), fill = "Experimental reads sens"),color="white") +
  scale_y_reverse(name="Percentage of reads") + 
  coord_cartesian(xlim = c(0, 80), ylim = c(83, 0)) + 
  scale_fill_manual(values = c("Experimental reads sens" = "#ff0000")) +
  #geom_freqpoly(data = PitsE, binwidth = 3,aes(x=Percentage_T,y=after_stat(count / tapply(count, group, sum)[group] * -100)*0, group=Sequence_Size,colour = factor(Sequence_Size)))+
  theme_classic() +
  scale_x_continuous(name = "Percentage of T",position = "top") +
  labs(color='Reads size',fill ='') +
  #scale_color_manual(values = oranges2) +
  #Descriptions on the figure  
  annotate("text", x = 25, y = 50, label = paste("Sens :", nrow(Pitssens)), color = "#ff0000",size = 7)+
  theme( legend.title = element_text(size = 10),legend.position = "none",
         legend.text = element_text( size = 10), axis.title.y = element_blank(),axis.title.x = element_blank(),
         axis.text.y = element_text(size = 18))

PitsPlot <- (PitsTop / PitsBottom)+
  plot_layout(heights = c(1, 1)) & 
  theme(plot.margin = margin(t = 0, b = 0))

ggsave(
  "Pits.pdf",
  PitsPlot,
  width = 25,
  height = 22,
  units = "cm",
)
```
